# Knocking out *ARL13B* completely abolishes primary ciliogenesis in cell lines

**DOI:** 10.1101/2025.03.17.643595

**Authors:** Divyanshu Mahajan, Hui Min Chia, Lei Lu

## Abstract

Small G protein ARL13B localizes to the cilium and plays essential roles in cilium biogenesis, organization, trafficking, and signaling. Here, we established multiple ARL13B knockout cell lines using the CRISPR/Cas9 system. Surprisingly, all our cell lines lost their cilia completely, in contrast to the reported short cilium and reduced ciliogenesis phenotype. We found that multiple regions of ARL13B are necessary for a complete rescue. Additionally, we found that ARL13B knockout cells also lost their response to SMO-mediated hedgehog stimulation. Our work demonstrates the critical requirement of ARL13B for ciliogenesis and hedgehog signaling, at least in cultured cells, and suggests that ARL13B plays a more crucial role in ciliary function than previously understood.

## Introduction

The primary cilium (hereafter cilium) is a microtubule-based plasma membrane protrusion that functions as the cell’s antenna. The cilium detects extracellular signals through cilium-localized receptors and triggers intracellular responses through various signaling pathways, such as hedgehog, Wnt, and TGFβ signal transduction pathways [1-3]. The critical role of the cilium in sensing and transducing signals for development and physiological function depends on its unique proteins, mutations of which can cause human disorders collectively termed ciliopathies [4, 5]. The ARF family small G protein ARL13B, the mutation of which causes the ciliopathy Joubert syndrome, has been recognized as one of the most crucial cilium regulators. ARL13B is highly enriched in the cilium, and such localization was mediated by the ciliary transport adaptor Rab8-TNPO1 or Tulp3 [6, 7]. Within the cilium, ARL13B recruits INPP5E to modulate the level of phosphoinositides [8-11]. ARL13B can also function as a guanine nucleotide exchange factor to activate another cilium-localized ARF family small G protein, ARL3, to transport lipidated cargos to the cilium [12-14].

Deletion of *ARL13B* in animals causes diverse cilium-associated physiological and cellular phenotypes [15-23]. For example, *ARL13B* null zebrafish (*scorpion* mutant) is characterized by cystic kidney and body curvature with significantly reduced cilium number and abnormal cilium organization [17]. *ARL13B* null mice (*hennin* mutant) have defective neural tubes, abnormal eyes, and axial polydactyly. At the molecular and cellular level, *hennin* mutant cells display reduced cilium length and biogenesis, abnormal cilium organization, and defective hedgehog signaling [16, 20]. With the advent of CRISPR/Cas9 technology, the *ARL13B* gene has been targeted to generate knockout (KO) cell lines. *ARL13B*-KO hTert-RPE1 (hereafter RPE1) and IMCD3 cells display short cilium with reduced ciliogenesis, consistent with observations in *ARL13B* null animals [9, 24]. In this work, we produced *ARL13B*-KO RPE1 and HEK293T cells and confirmed the KO of *ARL13B* by genotyping and Western blotting. Surprisingly, we found that cilia were completely lost in our *ARL13B*-KO cells. Concomitantly, the hedgehog signaling was abolished in these cells. Therefore, our data demonstrate that ARL13B might play a more critical role in cilium biogenesis and hedgehog signaling than previously recognized.

## Result

### *ARL13B*-KO RPE1 cells completely lack cilia

To study the cellular function of ARL13B, we employed CRISPR/Cas9-induced frameshift mutation to KO the *ARL13B* gene in RPE1 cells. We designed two sgRNAs, sg1 and 2, to target exons 1 and 5 of the human *ARL13B* gene, respectively (Fig. 1a). After lentivirus-mediated transduction of a single expression vector containing the sgRNA and Cas9, RPE1 cells were subjected to puromycin selection. We selected two stable cell lines, sg1#a and sg2#4, corresponding to sg1 and sg2, respectively. We sequenced the genomic DNA fragments containing the sgRNA target sites from sg1#a and sg2#4 cell lines. We found that the two cell lines have frameshift insertion or deletion of nucleotides in both alleles, which were expected to cause a premature translation termination of the transcribed mRNA and trigger the non-sense-mediated mRNA decay [25]. The open reading frames of frameshifted *ARL13B* in sg1#a and sg2#4 cell lines contain less than 13 and 169 AAs (Supplementary File 1), respectively, compared to wild type (WT) ARL13B, 428 AAs. Next, we assessed ARL13B protein levels in cell lysates prepared from sg1#a and sg2#4 cell lines using Western blotting. A single band at the expected molecular weight of ∼ 48 kDa was observed in parental RPE1 cells. In contrast, no band was observed in the two KO cell lines (Fig. 1b), supporting that sg1#a and sg2#4 are *ARL13B*-KO cells.

**Figure 1.**
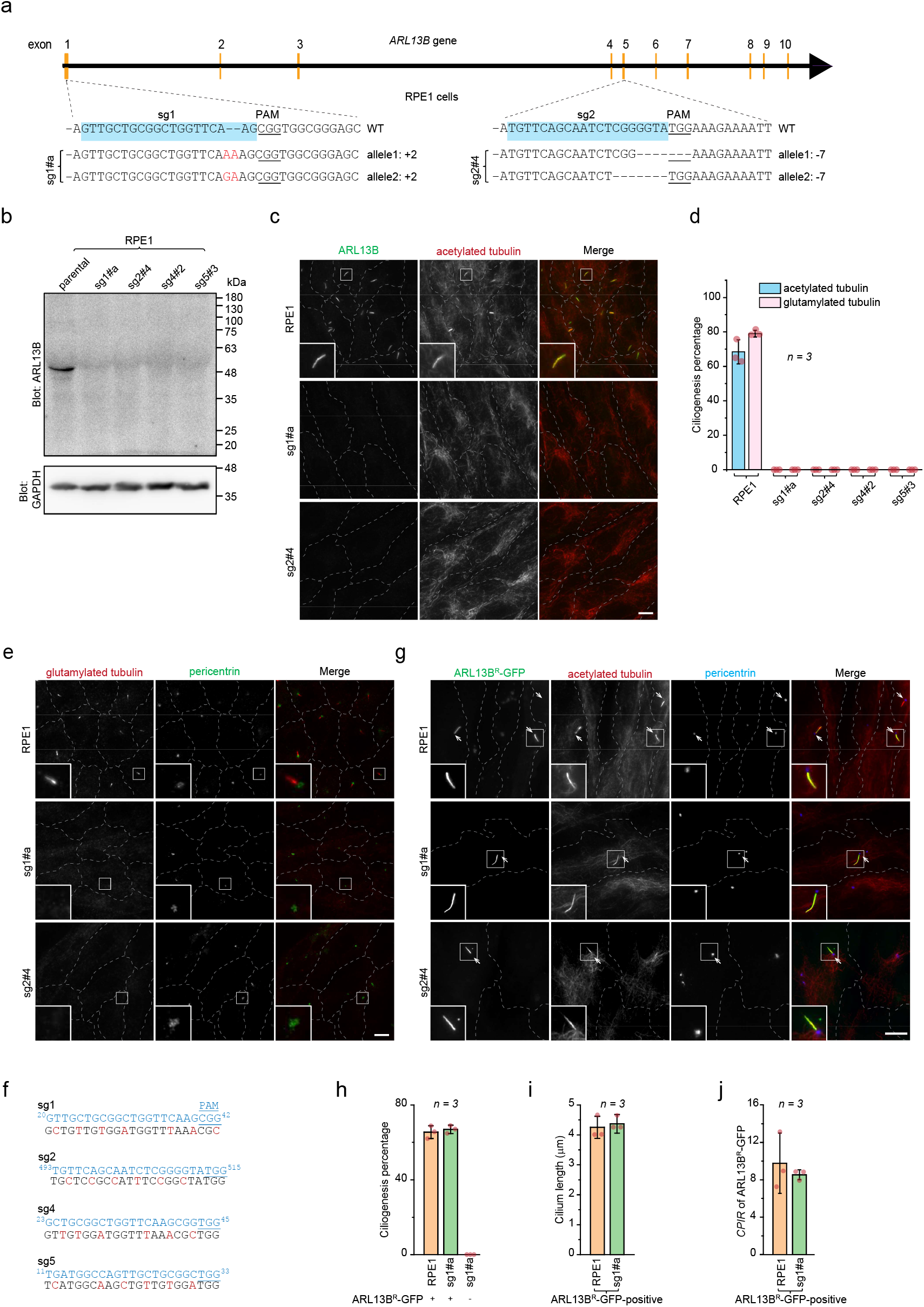
*ARL13B*-KO RPE1 cells have no cilia. **(a)** A schematic diagram of CRISPR/Cas9 strategy to make frameshift mutations in human *ARL13B* gene. The scheme of the exon organization of the human *ARL13B* gene is illustrated at the top. The middle shows parts of the WT genomic sequences of exon 1 and exon 5, where sequences of two sgRNAs, sg1 and sg2, are highlighted in light blue. Underlined sequences are protospacer-adjacent motifs or PAMs. The bottom shows genomic sequences at sgRNA target sites of two alleles in sg1#a and sg2#4 cell lines. Sequences are aligned with the corresponding WT ones. Inserted and deleted bases are colored red and indicated by “-”, respectively. “+2” and “-7” indicate insertion of 2 and deletion of 7 bases, respectively. **(b)** Western blotting of endogenous ARL13B in parental and KO RPE1 cells. Cell lysates from indicated cell lines were subjected to 12% SDS-PAGE followed by immunoblotting using anti-ARL13B and anti-glyceraldehyde 3-phosphate dehydrogenase (GAPDH) (loading control) antibodies. Molecular weight markers are labeled to the right of the immunoblot. **(c)** Fluorescence imaging of endogenous ARL13B and acetylated tubulin in parental and KO RPE1 cells. Indicated cells were fixed and co-immunostained by antibodies against indicated cilium markers before being subjected to wide-field microscopy. (**d**) Quantification of the ciliogenesis percentage in parental and KO RPE1 cells. Images acquired as described in (c) were analyzed. *n* = 3 independent experiments with ≥ 100 cells analyzed in each case. (**e**) Fluorescence imaging of endogenous glutamylated tubulin and pericentrin in parental and KO RPE1 cells, similar to (c). (**f**) Silent mutations in ARL13B^R^-GFP make it resistant to sgRNAs. Regions of the ARL13B^R^ coding sequence, corresponding to sg1, sg2, sg4, and sg5 (colored blue), are shown with introduced multiple silent mutations colored red. Protospacer-adjacent motifs, or PAMs, are underlined. Superscript numbers indicate base pair positions in the coding sequence. (**g**) ARL13B^R^-GFP can rescue the no-cilium phenotype of *ARL13B*-KO cell lines. Parental RPE1, sg1#a, and sg2#4 cells expressing ARL13B^R^-GFP were processed for immunofluorescence to label endogenous acetylated tubulin and pericentrin. Arrows indicate ciliated cells. In (c, e, and g), the boxed region is enlarged at the lower left corner, and the scale bar represents 10 μm. (**h-j**) ARL13B^R^-GFP expressing sg1#a cells exhibit normal cilia. Images acquired as described in (g) were used for quantification. Cilia were identified by acetylated tubulin. Panel h shows the ciliogenesis percentage in GFP-positive parental and sg1#a cells (indicated by “+”). It also displays the ciliogenesis percentage of GFP-negative sg1#a cells from the same coverslips (indicated by “-”). Quantification was from *n* = 3 independent experiments with ≥ 100 cells analyzed in each experiment. In (i), the cilium length was measured in ciliated GFP-positive cells. Quantification was from *n* = 3 independent experiments with ≥ 30 cells analyzed in each experiment. In (j), the *CPIR* of ARL13B^R^-GFP was quantified in ciliated GFP-positive cells. Quantification was from *n* = 3 independent experiments with ≥ 30 cells analyzed in each experiment. In (d, h-j), the error bar represents the *mean* ± *standard deviation*.

Serum starvation dramatically increased the ciliogenesis percentage of parental RPE1 cells from ∼ 20% to 70%, as revealed by immunostaining endogenous ARL13B, acetylated tubulin, or glutamylated tubulin (Fig. 1c,d), which are conventional cilium markers. Our immunofluorescence staining confirmed that the endogenous ARL13B level was undetectable in sg1#a and sg2#4 cells, as expected for *ARL13B*-KO. Surprisingly, we found no cilia in our two *ARL13B*-KO cell lines by the acetylated or glutamylated tubulin immunostaining. Our measured ciliogenesis percentages are 0.0 ± 0.0 % (*mean* ± *standard deviation* and same for the rest of the manuscript) (*n* = 3) in the two KO cell lines stained by the two tubulin cilium markers. Due to their possible short length and weak intensity, many acetylated or glutamylated tubulin-positive cilia might not be identified unambiguously against the cellular background in RPE1 cells. To facilitate the identification of cilia, we co-labeled the cilium and its base by immunostaining endogenous glutamylated tubulin and pericentrin [26] (Fig. 1e). In parental RPE1 cells, the cilium marked by glutamylated tubulin always extended from the cilium base marked by pericentrin-positive puncta. In contrast, we never observed cilia around the pericentrin puncta in sg1#a and sg2#4 cells. We also employed another cilium marker, fibrocystin ciliary targeting signal (CTS)-fused CD8a (CD8a-fCTS) [27] (Supplementary Fig. 1a, b), and we could not find cilia in our *ARL13B*-KO cell lines either. In summary, our data shows that *ARL13B*-KO RPE1 cells completely lack cilia.

To confirm that the no-cilium phenotype is not an off-target effect of gene editing, we performed rescue experiments by exogenously expressing CRISPR/Cas9-resistant ARL13B coding sequence (ARL13B^R^). ARL13B^R^ was constructed by making multiple silent point mutations within sg1 and sg2 target sites in the coding sequence of ARL13B (Fig. 1f). After the lentivirus-transduction of ARL13B^R^-GFP, sg1#a and sg2#4 cells were subjected to serum starvation and immunostaining of endogenous acetylated tubulin (Fig. 1g). Among GFP-positive cells, we observed 65 ± 3 % (*n* = 3) and 67 ± 2 % (*n* = 3), respectively, had cilia (Fig. 1h; Supplementary Fig. 1c). The ciliogenesis percentages of rescued cells had no significant difference from that of parental RPE1 cells (*p* = 0.001 by unpaired and two-tailed *t*-test). We previously introduced the cilium to plasma membrane intensity ratio (*CPIR*) as an expression level-independent metric that quantitatively measures the cilium localization [27]. We found that the *CPIR*s of ARL13B^R^-GFP and cilium lengths in rescued cells had no significant difference from those of parental RPE1 cells (Fig. 1i, j; Supplementary Fig. 1d, e). Therefore, our study demonstrates that ARL13B^R^-GFP can rescue the no-cilium phenotype in *ARL13B*-KO cells, suggesting that the no-cilium phenotype should be specific to *ARL13B*-KO.

### No cilia could be found in *ARL13B*-KO HEK293T cells either

We employed HEK293T cell lines to perform similar KO and characterization experiments to confirm our findings in another cell line. We used the same sgRNAs, sg1 and sg2, to generate *ARL13B*-KO cell lines, sg1#5 and sg2#5, respectively. Their genotypes confirmed the frameshift mutations in both alleles (Fig. 2a). By immunofluorescence imaging and Western blotting, we verified that ARL13B protein levels were undetectable in both cell lines (Fig. 2b, c). After serum starvation, we observed 43 ± 4 % (*n* = 3) parental HEK293T cells acquired acetylated or glutamylated tubulin-positive cilia (Fig. 2d, e). However, under identical treatment, we found no cilia in our two *ARL13B*-KO HEK293T cell lines, in which all ciliogenesis percentages were measured to be 0.0 ± 0.0 % (*n* = 3). We also demonstrated that the no-cilium phenotype in our *ARL13B*-KO HEK293T cell lines could be rescued by exogenously expressing ARL13B^R^-GFP (Supplementary Fig. 2). Therefore, our data confirms the no-cilium phenotype upon *ARL13B*-KO in HEK293T cells and suggests that such an observation might apply to multiple types of cells.

**Figure 2.**
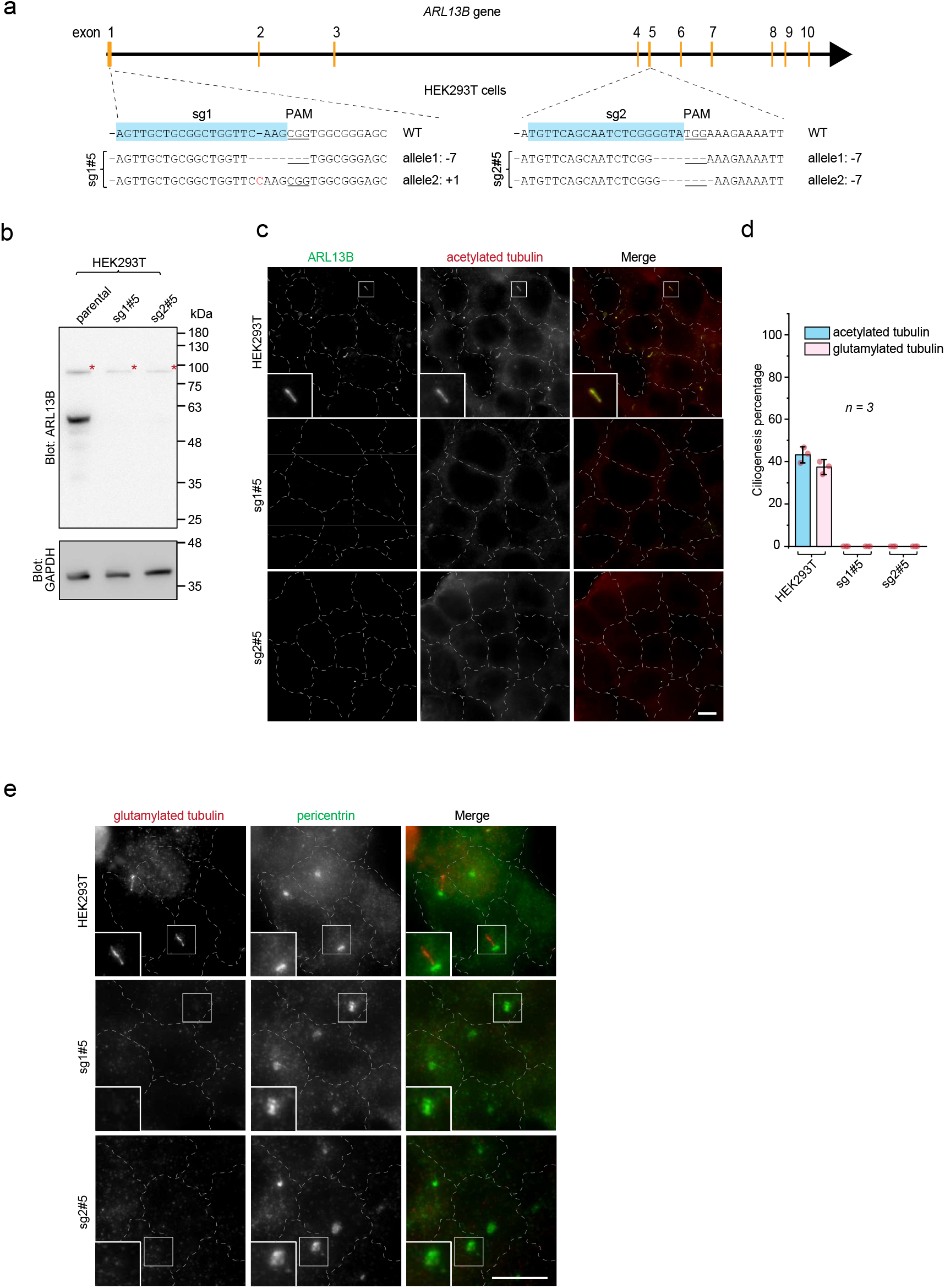
*ARL13B*-KO HEK293T cells have no cilia. **(a)** The genotypes of *ARL13B*-KO HEK293T cell lines, sg1#5 and sg2#5. See the legend of Figure 1a. (**b-e**) Endogenous ARL13B is undetectable by Western blot and immunofluorescence imaging, and no cilia could be found in sg1#5 and sg2#5 cells. In (**b**), anti-GAPDH was used for loading control. Non-specific bands are indicated by asterisks (*). Molecular weight markers are labeled to the right of the immunoblot. See the legend of Figure 1b. (**c**) Fluorescence imaging of endogenous ARL13B and acetylated tubulin in parental and KO HEK293T cells. Cells were fixed and co-immunostained by antibodies against indicated cilium markers before being subjected to wide-field microscopy. (**d**) Quantification of the ciliogenesis percentage in parental and KO HEK293T cells. Images acquired as described in (c) were analyzed. *n* = 3 independent experiments with ≥ 100 cells analyzed in each case. The error bar represents the *mean* ± *standard deviation*. (**e**) Fluorescence imaging of endogenous glutamylated tubulin and pericentrin in parental and KO HEK293T cells, similar to (c). In (c and e), the boxed region is enlarged at the lower left corner, and the scale bar represents 10 μm.

### *ARL13B*-KO by previously reported sgRNAs also displayed the no-cilium phenotype

The no-cilium phenotype we observed in two *ARL13B*-KO cell lines differs from previous studies, such as *ARL13B* null mice (*hennin*) [16, 20] and zebrafish (*scorpion*) [17]. It is especially in stark contrast to a similar work by Nozaki *et al*., who produced *ARL13B*-KO RPE1 cells using CRISPR/Cas9 technique [9]. Nozaki *et al*. reported that cilia were still clearly present in *ARL13B*-KO cells but with a reduced length. To resolve this discrepancy, we adopted the same two sgRNAs they designed, sg4 and sg5, corresponding to sgRNA #1 and #2 in their study, to knock out *ARL13B* gene in the same RPE1 cells. Our effort resulted in two clonal cell lines, sg4#2 and sg5#3, respectively (Fig. 3a). Their frameshift insertion and deletion genotypes and undetectable expression levels of ARL13B in Western blot and immunofluorescence confirmed their ARL13B null phenotypes in both cell lines (Fig. 1b; Fig. 3b). Importantly, consistent with our sg1#a, sg2#4, sg1#5, and sg2#5 cells, we still observed a complete loss of cilia by immunostaining of endogenous acetylated and glutamylated tubulin with a ciliogenesis percentage of 0.0 ± 0.0 % (*n* = 3) for both cell lines (Fig. 3b, c). Similarly, we also demonstrated that the no-cilium phenotype was rescued by exogenously expressing ARL13B^R^-GFP (Fig. 3d).

**Figure 3.**
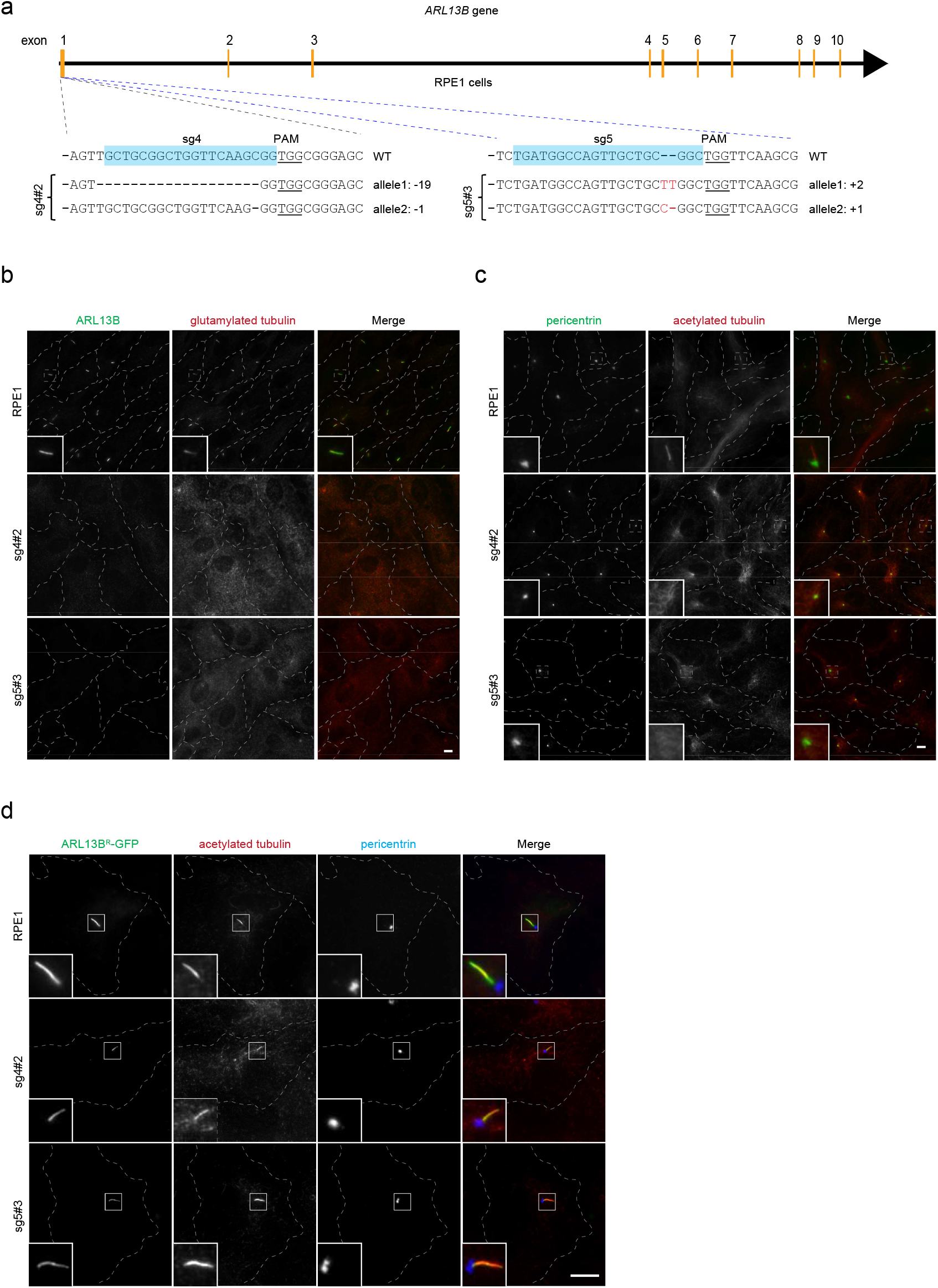
*ARL13B*-KO RPE1 cells prepared using previously reported sgRNAs have no cilia either. (**a**) The genotypes of *ARL13B*-KO RPE1 cell lines, sg4#2 and sg5#3, using previously published sgRNAs. See the legend of Figure 1a. (**b-c**) Endogenous ARL13B is undetectable by immunofluorescence imaging and no cilia could be found in sg4#2 and sg5#3 cells. See the legend of Figure 1c, e (**d**) Exogenously expressed ARL13B^R^-GFP can rescue the no-cilium phenotype of sg4#2 and sg5#3 cells. See the legend of Figure 1g. In (b-d), the boxed region is enlarged at the lower left corner, and the scale bar represents 10 μm.

Gene KOs generated by the CRISPR/Cas9-induced frameshift mutation can sometimes still express residual target protein, resulting in a weak phenotype [28-30]. Since our KO cells exhibit a much stronger phenotype, no-cilium, than that reported by Nozaki *et al*. and others, which is reduced cilium length and ciliogenesis percentage [9, 16, 17], we believe our KO cells likely represent a more complete deletion of the *ARL13B* gene (see Discussion). In summary, using two sgRNAs in two cell lines and replicating a previous study, we demonstrated that *ARL13B*-KO cells lack cilia entirely rather than merely exhibiting reduced cilium length and biogenesis.

### Multiple ARL13B regions or domains are essential for its ciliogenesis function

To investigate the roles of different ARL13B regions or domains in ciliogenesis, we assessed the rescue of the no-cilium phenotype of sg1#a cells using a series of ARL13B mutants. ARL13B consists of the following regions or domains from the N to the C-terminus: an α-helix region with palmitoylation lipid anchorage sites (PAL), a GTPase domain, a coiled-coil region (CC), an intrinsically disordered region (IDR), the RVEP motif, and a proline-rich region (PRR) (Fig 4a). The N-terminal PAL contains two palmitoylation sites at C8 and C9, which enable the membrane anchorage of ARL13B. Mutation of both palmitoylation sites from cysteine to serine (C8S/C9S) in PAL abolishes the cilium localization of ARL13B [6, 24, 31, 32]. The GTPase domain is essential for the interaction with INPP5E and can act as a guanine nucleotide exchange factor for ARL3 [8, 9, 12-14]. According to the structural and functional analysis of Ras superfamily (including ARF family) small G proteins, the T35N mutation of the GTPase domain’s G1 box is expected to lock ARL13B in a GDP-binding state [33-35]. The cellular function of the CC remains elusive, but it is essential for the cilium localization of ARL13B and INPP5E interaction [6, 9]. While the function of the IDR is still unclear, the PRR is known to interact with the IFT-B complex and INPP5E [9]. Although both IDR and PRR are dispensable for the cilium localization of ARL13B, the RVEP motif, a stretch of 17 AAs located between the IDR and PRR, serves as the CTS of ARL13B by interacting with ciliary transport adaptors, TNPO1 and Rab8 [6].

**Figure 4.**
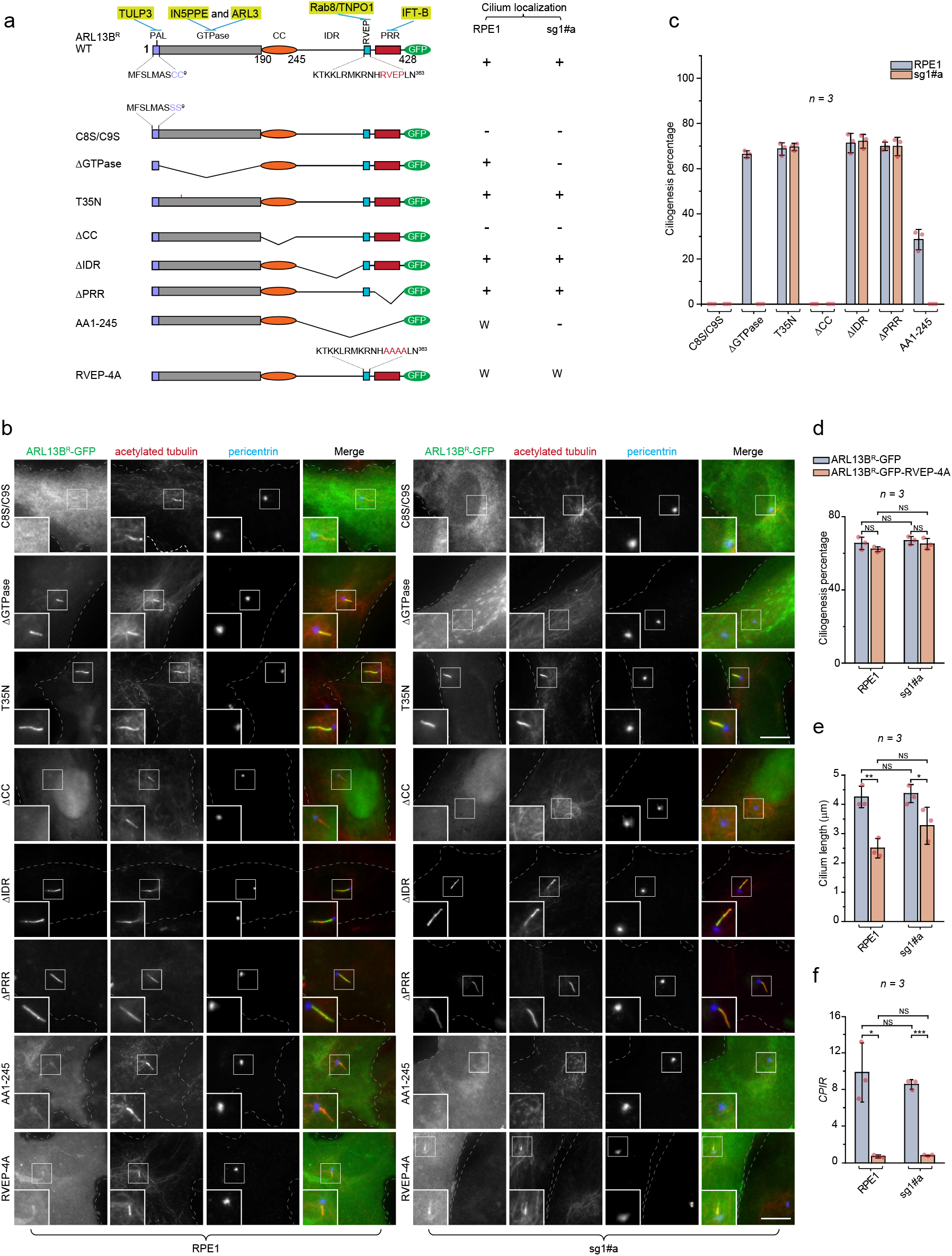
Multiple ARL13B regions or domains are required for ciliogenesis. (**a**) The schematic diagram illustrating the organization of WT and mutant ARL13B, which were constructed in the background of ARL13B^R^-GFP. The top shows the WT ARL13B. Different functional regions and domains are indicated with AA positions. Interacting proteins previously reported are labeled above the corresponding regions or domains and shaded green. The AA sequences of PAL and RVEP motifs are shown below. ARL13B mutants with point mutations or truncations are illustrated below the WT. On the right is the qualitative summary of the cilium localization of corresponding GFP-tagged WT and mutant ARL13B (see below for details). “+” indicates a positive cilium localization; “-” indicates a negative cilium localization or no ciliogenesis; “W” indicates a weak cilium localization. (**b**) The cellular distribution of GFP-tagged WT and mutant ARL13B^R^-GFP. Parental and sg1#a RPE1 cells expressing GFP-tagged WT and mutant ARL13B^R^-GFP were immunostained for indicated proteins. The boxed region is enlarged at the lower left corner, and the scale bar represents 10 μm. (**c**) The cilium localization percentage (Parental RPE1 cells) and cilium biogenesis percentage (sg1#a RPE1 cells) of GFP-positive cells. The images acquired as described in (b) were analyzed. *n* = 3 independent experiments with ≥ 100 cells analyzed in each. (**d-f**) The RVEP motif is non-essential for ciliogenesis but essential for the cilium length and the cilium enrichment of ARL13B^R^-GFP in *ARL13B*-KO cells. Images acquired as described in (b) were quantified. GFP-positive cells were analyzed for the ciliogenesis percentage (**d**), the cilium length (**e**), and the *CPIR* (**f**) of ARL13B^R^-GFP. See legend of Figure 1h-j. *p*-values are from unpaired, two-tailed *t*-tests. Not significant or NS, *p* > 0.05; *, *p* ≤ 0.05; **, *p* ≤ 0.005; ***, *p* ≤ 0.0005. In (c, d-f), the error bar represents the *mean* ± *standard deviation*.

We prepared the following mutants in the background of ARL13B^R^-GFP: C8S/C9S, ΔGTPase (deletion of GTPase domain), ΔCC (deletion of CC), ΔIDR (deletion of IDR), RVEP-4A (mutation of RVEP to AAAA), ΔPRR (deletion of PRR), and AA1-245 (deletion of AAs from 246 to 428) (Fig. 4a). In parental RPE1 cells, we demonstrated that C8S/C9S and ΔCC do not localize to the cilium, while ΔGTPase, ΔIDR, and ΔPRR show robust cilium localization and RVEP-4A and AA1-245 exhibit weak cilium localization (Fig. 4a; Fig. 4b, parental RPE1 cells), consistent with our previous study [6]. Additionally, we observed that T35N has a robust cilium localization (Fig. 4a; Fig. 4b, parental RPE1 cells).

Next, we tested these ARL13B mutants for their ability to rescue the no-cilium phenotype of sg1#a cells. Following lentivirus-mediated transduction and serum starvation, we analyzed ciliogenesis by immunostaining endogenous acetylated tubulin and pericentrin (Fig. 4b, c). We found that T35N, ΔIDR, and ΔPRR successfully rescued the no-cilium phenotype with a ciliogenesis percentage of ≥ 70 %, showing no significant difference from ARL13B^R^-GFP (*p* > 0.05) (Fig. 4c). In contrast, no cilia were found in C8S/C9S, ΔGTPase, ΔCC, and AA1-245 expressing sg1#a cells, all exhibiting a ciliogenesis percentage of 0.0 ± 0.0 % (*n* = 3) (Fig. 4c).

Interestingly, although RVEP-4A-expressing sg1#a cells did not show a significant difference (p > 0.05) from WT-expressing cells in terms of the ciliogenesis percentage, 65 ± 3 % vs. 67 ± 2 % (*n* = 3) (Fig. 4d), the resulting cilia were significantly (*p* < 0.05) shorter than WT, measuring 3.3 ± 0.6 vs 4.4 ± 0.3 μm, respectively (*n* = 3)(Fig. 4e). Our observation of the RVEP-4A mutant is consistent with a previous study where ARL13B with V358A mutation did not rescue the cilium length in *ARL13B* null *hennin* mouse embryonic fibroblasts [36]. The partial rescue phenotype might be due to the insufficient amount of cilium-localized ARL13B, as revealed by its significantly (*p* < 0.05) lower *CPIR* than WT in parental RPE1 cells, at 0.61 ± 0.04 vs. 8.5 ± 0.5 (*n* = 3), respectively (Fig. 4f). This observation is consistent with previous reports showing that the ciliary level of ARL13B regulates cilium length [20, 37]. In parental RPE1 cells, overexpression of the RVEP-4A mutant also significantly reduced cilium length compared to WT ARL13B, with lengths of 2.5 ± 0.3 µm vs. 4.3 ± 0.4 µm, respectively (*p* < 0.005, *n* = 3) (Fig. 4e). It is possible that overexpression of the RVEP-4A mutant, as demonstrated by our quantitative immunofluorescence imaging (Supplementary Fig. 3), interferes with the ciliary targeting of endogenous ARL13B in a dominant-negative manner, thereby contributing to the short cilium phenotype.

In summary, we demonstrated that our *ARL13B-KO* cells provided an excellent tool to test the ciliogenesis role of ARL13B domains, motifs, or mutations, as evidenced by the clear contrast with the no-cilium phenotype during the rescue. Our extensive mutagenesis demonstrated that the ciliogenesis function of ARL13B requires the N-terminal palmitoylation, the GTPase domain, and the CC. We found that cilium localization of ARL13B mutants is necessary but insufficient for ciliogenesis, as shown by ΔGTPase and AA1-245. Furthermore, our data on the RVEP-4A mutant suggests that cilium formation requires a minimal amount of ciliary ARL13B, while the dose of cilium-localized ARL13B might be a limiting factor for cilium length.

### Joubert syndrome mutation, R200C, abolishes the ciliogenesis function of ARL13B

We focused on the three point mutations in the coding sequence of the *ARL13B* gene, which have been associated with the Joubert syndrome [38, 39]. Among them, R79Q, and Y86C are in the GTPase domain, while R200C is in the CC. To characterize the impact of these point mutations on the ciliogenesis function of ARL13B, we made corresponding mutations in the background of ARL13B^R^-GFP (Fig. 5a). First, we examined their cilium localization in parental RPE1 cells, similar to our approach with other mutants. We found that R79Q and Y86C had robust cilium localization comparable to WT (Fig. 5b). The cilium localization of R200C was significantly weaker than that of WT (Fig. 5b). Next, we tested if ARL13B with these Joubert syndrome point mutations can rescue the no-cilium phenotype of sg1#a cells (Fig. 5c). Following lentivirus-mediated transduction, serum starvation, and immunostaining, we analyzed the ciliogenesis of these mutants. R79Q and Y86C displayed ciliogenesis percentages of 71 ± 3 % (*n* = 3) and 73 ± 1 % (*n* = 3) in sg1#a cells (Fig. 5d), respectively, similar to WT (Fig. 1h). In contrast, no cilia were observed in R200C-expressing sg1#a cells, which exhibited a ciliogenesis percentage of 0.0 ± 0.0 % (*n* = 3) (Fig. 5d). In summary, our data revealed that R200C completely abolishes the ciliogenesis function of ARL13B, highlighting the essential role of the CC in ciliogenesis and suggesting a cellular mechanism of R200C mutation for the Joubert syndrome pathology. In contrast, the R79Q and Y86C mutations do not appear to impair ciliogenesis, implying that their cellular effects might act downstream of cilium formation.

**Figure 5.**
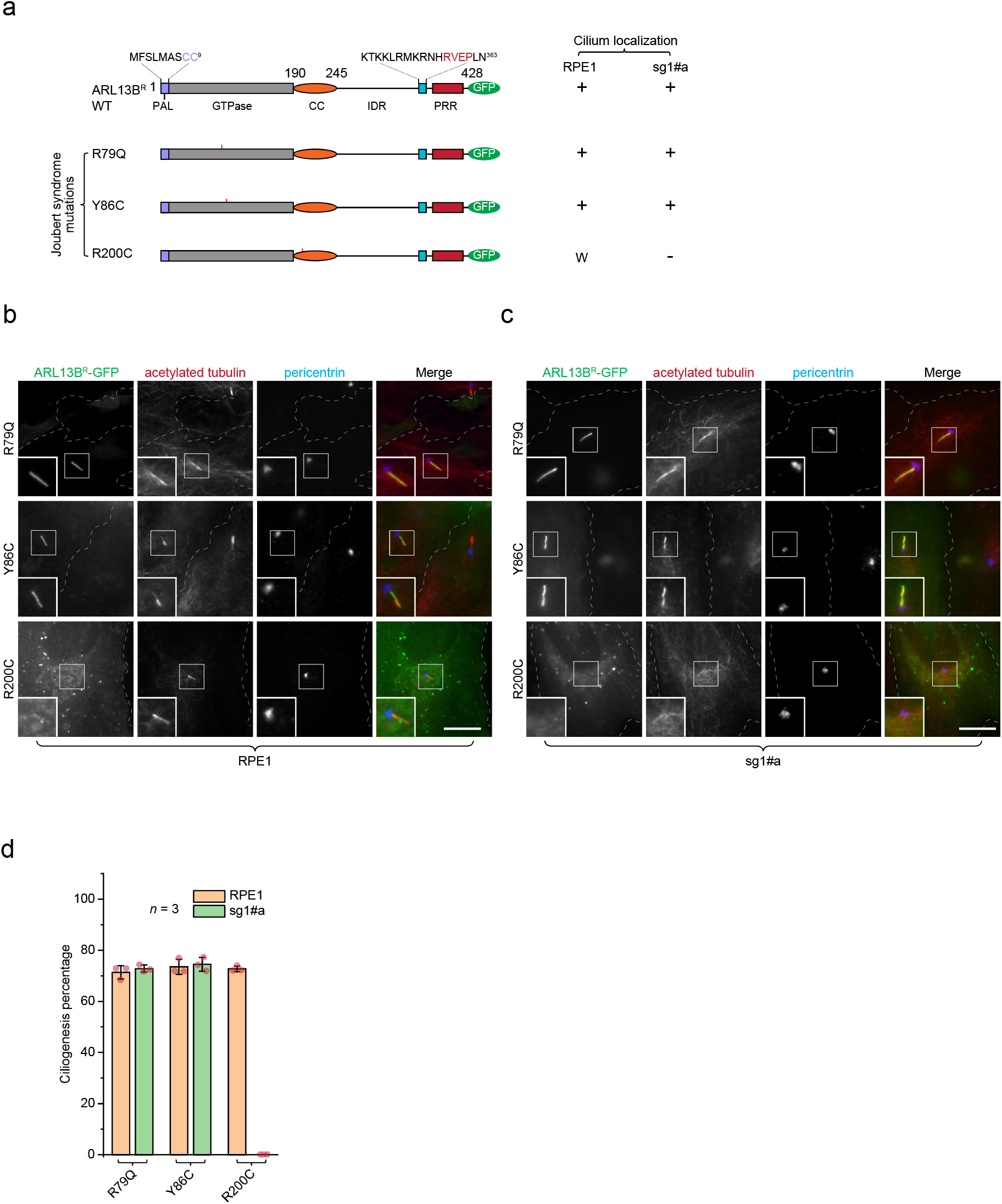
Joubert syndrome mutation R200C, but not R79Q and Y86C, is essential for the ciliogenesis function of ARL13B. (**a**) The schematic diagram illustrates the organization of ARL13B and its three Joubert syndrome mutants. See the legend of Figure 4a. (**b-c**) The cellular distribution, cilium localization, and ciliogenesis percentage of GFP-tagged Joubert syndrome mutant ARL13B^R^. The boxed region is enlarged at the lower left corner, and the scale bar represents 10 μm. See the legend of Figure 4b, c. (**d**) The images acquired as described in (b-c) were analyzed. *n* = 3 independent experiments with ≥ 30 cells analyzed in each. The error bar represents the *mean* ± *standard deviation*.

### Hedgehog signaling is abolished in *ARL13B*-KO cells

It is known that the hedgehog signaling requires the cilium [40]. The stimulation of hedgehog recruits SMO to the cilium and upregulates the transcription of hedgehog signaling target genes, such as *SMO, PTCH1*, and *GLI1*. Previous studies established that the loss of ARL13B disrupts the hedgehog signaling in vertebrates such as mouse (*hennin* mutant) and zebrafish (*scorpion* mutant) [16, 17], which have the phenotypes of reduced cilia. Therefore, we want to investigate the impact of a complete cilium loss on the hedgehog signaling pathway in our *ARL13B*-KO cells by assessing the cilium localization of SMO and measuring the transcription of *SMO, PTCH1*, and *GLI1* genes.

In serum-starved parental RPE1 cells, we immunostained the endogenous ARL13B and SMO. We observed that only 11 ± 3 % (*n* = 4) of ciliated cells had the cilium localization of SMO (Fig. 6a, b control). In comparison, after 24 h treatment with 0.5 μM SAG (SMO agonist), a small molecule that directly activates SMO, the cilium localization of SMO significantly (*p* < 0.005) increased to 75 ± 5 % (*n* = 4) in ciliated cells (Fig. 6a, b). Using qPCR, we found that SAG treatment after serum starvation increased the normalized transcription levels of *SMO, PTCH1*, and *GLI1* genes to 2.3 ± 0.9, 1.6 ± 0.3, and 4.5 ± 3.6, respectively, (*n* = 5, normalized by control), respectively (Fig. 6c). Our results are consistent with what we know about the SMO-activated hedgehog signaling, thereby validating our assay setup.

**Figure 6.**
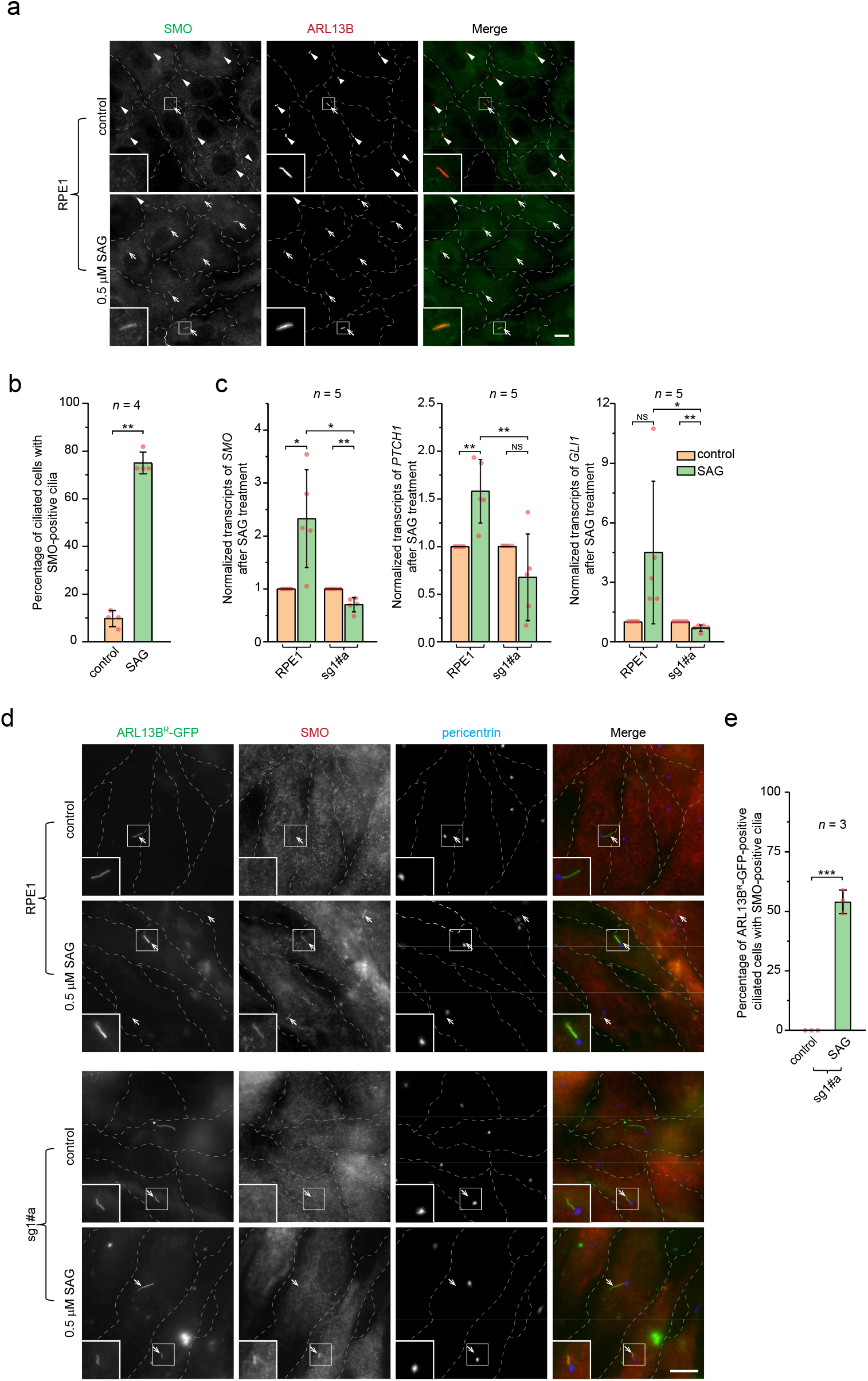
Hedgehog signaling is abolished in *ARL13B*-KO cells. (**a**) SAG treatment increased the cilium localization of SMO in RPE1 cells. RPE1 cells treated with or without (control) 0.5 μM SAG for 24 hours were fixed and processed for immunofluorescence labeling of endogenous SMO and ARL13B. Arrows and arrowheads indicate SMO-positive and -negative cilia, respectively. (**b**) Percentage of ciliated cells with SMO-positive cilia. Images acquired as described in (a) were analyzed. *n* = 4 independent experiments. (**c**) Normalized transcription levels of *SMO, PTCH1*, and *GLI1* genes in parental RPE1 or sg1#a cells with or without (control) 0.5 μM SAG treatment for 24 h. The transcription was quantified by qPCR and normalized by the corresponding control. *n* = 5 independent experiments. (**d-e**) Exogenously expressed ARL13B^R^-GFP can rescue the SAG-induced cilium localization of SMO in *ARL13B*-KO cells. Parental RPE1 or sg1#a cells expressing ARL13B^R^-GFP were treated with or without (control) 0.5 μM SAG for 24 h. Cells were subsequently fixed and processed for immunofluorescence labeling of endogenous SMO and pericentrin (d). Arrows indicate ciliated cells. The percentage of ARL13B^R^-GFP expressing ciliated sg1#a cells with SMO-positive cilia is quantified in (e). *n* = 3 independent experiments with ≥ 100 cells analyzed in each. In (a, d), scale bar, 10 µm. In (b, c, e), the error bar represents the *mean ± standard deviation*, and *p*-values were from unpaired, two-tailed *t*-tests. NS, *p* > 0.05; *, *p* ≤ 0.05; ** *p* ≤ 0.005; *** *p* ≤ 0.0005.

Since *ARL13B*-KO sg1#a cells have no cilia, hedgehog signaling components, including SMO, could not localize to the cilium. To assess the hedgehog signaling upon the KO of *ARL13B*, we quantified the hedgehog-dependent transcription of *SMO, PTCH1*, and *GLI1* genes. First, we found no significant difference in the transcript levels of *SMO, PTCH1*, and *GLI1* genes between parental RPE1 and sg1#a cells (Supplementary Fig. 4). Next, we treated sg1#a and parental RPE1 cells in parallel with serum-starvation followed by SAG, as described in Figure 6c. In contrast to the parental RPE1 cells, we found that SAG treatment did not increase the relative transcription levels of *SMO, PTCH1*, and *GLI1* genes; instead, it slightly reduced their levels to 0.7 ± 0.1, 0.7 ± 0.5, and 0.7 ± 0.2, respectively, (*n* = 5, normalized by untreated). Among these, the reductions in *SMO* and *GLI1* transcription were statistically significant, whereas the change in *PTCH1* was not. The biological significance of the decreased transcription of *SMO* and *GLI1* remains unclear and warrants further investigation.

The exogenous expression of ARL13B^R^-GFP rescued the SAG-induced cilium localization of SMO in sg1#a cells, as we observed an increase in the cilium localization of SMO among GFP-positive ciliated cells from 0 % to 56 ± 5 % (*n* = 3) following SAG treatment (Fig. 6d,e), demonstrating that the observed effect on hedgehog signaling is specific to *ARL13B*-KO. In summary, our results revealed that hedgehog signaling is severely blunted upon the KO of *ARL13B*, suggesting that *ARL13B*-KO cells might become irresponsive to hedgehog.

## Discussion

*ARL13B* null mice and zebrafish have been reported previously to display a reduced but still appreciable ciliogenesis percentage, cilium length, and hedgehog signaling. Similarly, *ARL13B*-KO RPE1 and IMCD3 cells have been shown to form cilia, albeit with reduced length [9, 24]. However, our *ARL13B*-KO cells, generated using four sgRNAs in two cell types, RPE1 and HEK293T, consistently exhibited much more extreme phenotypes — both cilia and hedgehog signaling appear to be completely lost, which is in stark contrast to many previous studies. Despite using the same cell type (RPE1) and technique (CRISPR/Cas9), our findings differ significantly from those of Nozaki et al., as we observed a non-cilium phenotype even after adopting the same sgRNAs. However, our results are supported by those of Shi *et al*., who reported cilia loss in *ARL13B*-KO glioma cell lines generated using CRISPR/Cas9 [41], and by Li *et al*., who observed the absence of cilia in specific kidney cell types following *ARL13B* deletion via the Cre-LoxP system in mice [21].

It has recently been reported that KO clones generated by CRISPR/Cas9-induced frameshift often express a residual amount of the target protein and, therefore, can produce much milder phenotypes than expected [28-30]. The low expression of the target protein in these “pseudo-KO” clones could result from multiple mechanisms, such as alternative splicing or usage of the downstream start codon of the target mRNA. Thus, it is tempting to speculate that *ARL13B*-KO clones and null mutants generated by Nozaki *et al*. and other labs could be such cases with a low level of ARL13B protein but still sufficient to support ciliogenesis. Hence, their phenotypes are weaker than ours. Supporting this hypothesis, we found that reducing the ciliary localization of ARL13B by mutating its CTS, the RVEP motif, did not affect ciliogenesis but significantly shortened the cilium length (Fig. 4e). To confirm this hypothesis, we propose three possible experiments using these “pseudo-KO” cells. First, the residual amount of the target protein can be enriched and subjected to a more sensitive Western blotting assay. Second, reverse transcription PCR can be employed to detect alternative splicing events of the *ARL13B* mRNA. Lastly, further knockdown of ARL13B by RNAi in these cells should produce no-cilium phenotypes similar to ours.

Our data revealed the crucial requirement of ARL13B in ciliogenesis. Our rescue experiments in *ARL13B*-KO cells using ARL13B mutants further suggest the roles of different ARL13B regions and domains in ciliogenesis. The palmitoylation membrane anchorage, the GTPase domain, and the CC, but not the IDR and PRR, are required for ciliogenesis. In addition, the guanine nucleotide exchange of ARL13B might not be required, as suggested by the GDP-locked mutation, T35N. The RVEP motif regulates the ciliary localization of ARL13B, which is essential for cilium growth. Mutation of this motif, as in RVEP-4A, impairs ARL13B targeting to the cilium and results in a short-cilium phenotype (Fig. 4e). Additionally, we examined three Joubert syndrome point mutations and found that R200C, but not R79Q and Y86C, abolished the ciliogenesis. Therefore, our data also help us to understand the pathogenesis of Joubert syndrome with R200C mutation.

## Material and methods

### DNA plasmids

Please see Supplementary Table 1 for all the plasmids used in this study. pLenti-CRISPR v2 was a gift from Feng Zhang (Addgene plasmid # 52961; http://n2t.net/addgene:52961; RRID: Addgene_52961).

### Antibodies

Mouse monoclonal antibodies (mAbs) against acetylated α-tubulin (Merck, #T6793, 1:1000 for immunofluorescence or IF), polyglutamylated tubulin (Adipogen, #GT335, 1:1000 for IF), CD8a (Developmental Studies Hybridoma Bank, clone OKT8, 1:200 for IF), GAPDH (Santa Cruz, #sc25778, 1:1000 for WB), and SMO (Santa Cruz, #sc-166685) Rabbit polyclonal antibodies (pAbs) against ARL13B (in-house made; 1:1000 for IF)[27], ARL13B (ProteinTech, #17711-AP, 1:1000 for IF and 1:3000 for WB), and pericentrin (Abcam, #ab4448, 1:1000 for IF). Horseradish peroxidase (HRP) -conjugated goat anti-mouse and anti-rabbit IgG antibodies were purchased from Bio-Rad. Alexa Fluor-conjugated goat anti-mouse and anti-rabbit IgG antibodies (1:500 for IF) were purchased from Invitrogen.

### Cell culture and transfection

hTert-RPE1 and HEK293T cells were from the American Type Culture Collection. 293FT cells were from Thermo Fisher Scientific. RPE1 cells were cultured in Dulbecco’s Modified Eagle’s Medium (DMEM) and Ham’s F12 mixture medium supplemented with 10% fetal bovine serum (FBS). HEK293T cells were cultured in DMEM supplemented with 10% FBS. 293FT were cultured in DMEM supplemented with 10% FBS and 500 μg/ml Geneticin™ (Thermo Fisher Scientific, #10131035). Transfection of plasmid DNA was performed using polyethylenimine (Polysciences) or Lipofectamine 2000 (Thermo Fisher Scientific, #11668019), according to the manufacturer’s protocol. Ciliogenesis was induced by incubating cells in the corresponding medium without the FBS for 48 h (serum starvation).

### Lentivirus-transduced KO and rescue of *ARL13B*

sgRNA cloned in pLenti-CRISPRV2 vector or ARL13B^R^ WT and mutants cloned in pLVX-puro vector were transiently transfected with packaging plasmids pLP1, pLP2, and pLP/VSVG (Thermo Fisher Scientific) to 293FT cells using Lipofectamine 2000 according to the manufacturer’s protocol. Lentiviruses were harvested 36 and 60 h post-transfection, filtered through a 0.45 µm syringe-driven filter (Sartorius), and immediately used to infect WT RPE1, HEK293T, or *ARL13B*-KO cell lines.

To make *ARL13B*-KO cell lines, RPE1 or HEK293T cells seeded in a 6-well plate were infected twice every 24 h with filtered lentiviruses using 8 μg/ml hexadimethrine bromide (Sigma-Aldrich, #H9268). 24 h after the second infection, the cells were then selected in media supplemented with 8 and 1 μg/ml puromycin (Sigma-Aldrich, #P8833) for 72 h for RPE1 and HEK293T cells, respectively. Subsequently, highly diluted cells were seeded into a 96-well plate so that each well had at most one colony. Colonies were harvested, expanded, and characterized for *ARL13B*-KO using immunofluorescence and Western blotting. At least one clonally selected cell line for each sgRNA was selected and sequenced around the sgRNA target site to confirm frameshift mutations in both.

In rescue experiments, *ARL13B*-KO cells seeded in a 6-well plate were infected twice every 24 h with filtered lentiviruses using 8 μg/ml hexadimethrine bromide. The cells were then seeded into a 24-well plate containing coverslips for immunofluorescence followed by serum starvation for 48 h to induce ciliogenesis before immunolabeling. For the hedgehog signaling assay, 24 h treatment with 0.5 μM SAG (Abcam, #ab142160) was performed before immunolabeling.

### Western blot

Parental and *ARL13B-KO* cells seeded in a 6-well plate were lysed in the SDS sample buffer and resolved in SDS-PAGE. SDS-PAGE separated proteins were transferred to polyvinyl difluoride membrane (Bio-Rad), which was sequentially incubated with primary and HRP-conjugated secondary antibody. Western blot and molecular weight marker bands were acquired under the chemiluminescence and white-light imaging mode, respectively, using a cooled charge-coupled device of LAS-4000 (GE Healthcare Life Sciences). Molecular weights were manually assigned by aligning the two images. Uncropped blot images are presented in Supplementary Fig. 5.

### RT-qPCR for SAG-activated hedgehog signaling pathway

RPE1 and *ARL13B*-KO cells were cultured to confluence in a 6-well plate, followed by serum starvation for 48 h to induce ciliogenesis. After 48 h of starvation, the control sample was cultured for another 24 h in the serum starvation medium, and the SAG sample was subjected to 0.5 μM SAG. Total RNA was extracted using TRIzol™ Reagent (Invitrogen™, #15596025) and reverse-transcribed, primed by random nonamer primers using M-MuLV Reverse Transcriptase (NEB, #M0253), following the manufacturer’s standard protocol. The qPCR reactions were performed using the iTaq Universal SYBR Green Supermix (Bio-Rad, #1725121), in accordance with the manufacturer’s instructions. Gene expression data for β actin, *SMO, PTCH1*, and *GLI1* were acquired using Bio-Rad CFX Connect Real-Time PCR Detection System. The qPCR result of each gene was first divided by that of β actin and further normalized to that of control. The relative gene expression was calculated through 2^-ΔΔCT^ method. Primer designed for the target genes are listed in Supplementary Table 1.

### Immunolabeling and wide-field fluorescence microscopy

RPE1 and HEK293T cells were first seeded on No. 1.5 Φ 12-mm coverslips in a 24-well plate. Next, cells were serum starved for 48 h to induce ciliogenesis, fixed with 4% paraformaldehyde in PBS, and subsequently neutralized with ammonium chloride. Subsequently, the cells were incubated with primary antibodies diluted in the antibody dilution buffer, which consists of PBS supplemented with 5% FBS, 2% bovine serum albumin, and 0.1% Saponin (Sigma-Aldrich). After several washings, cells were incubated with secondary antibodies diluted in the antibody dilution buffer. The coverslips were mounted in Mowiol 4-88 (EMD Millipore) after extensive washing using PBS. Finally, cells were imaged under an inverted wide-field Olympus IX83 microscope system equipped with a Plan Apo oil objective lens (63×, NA 1.40), a motorized stage, a focus drift correction device, motorized filter cubes, a scientific complementary metal oxide semiconductor camera (Neo; Andor Technology), and a 200 W metal halide excitation light source (Lumen Pro 200; Prior Scientific). Dichroic mirrors and filters in filter turrets were optimized for Alexa Fluor 488 (or GFP), Alexa Fluor 594, and Alexa Fluor 680. The microscope system was controlled by MetaMorph software 7.8.3.0 (Molecular Devices, https://support.moleculardevices.com/s/article/MetaMorph-Software-installation-files).

### Image analysis for the ciliogenesis percentage, the cilium length, and the *CPIR*

Untransfected cells were serum-starved to induce ciliogenesis. They were subsequently fixed and subjected to immunofluorescence labeling for endogenous cilium markers before wide-field microscopy imaging. The ciliogenesis percentage was calculated by dividing the number of ciliated cells by the total number of analyzed cells. Parental or *ARL13B*-KO cells expressing ARL13B^R^-GFP WT and mutants were similarly processed and imaged. Their ciliogenesis percentages were calculated by dividing the number of ciliated GFP-positive cells by the total number of analyzed GFP-positive cells. The cilium length and the *CPIR* of ARL13B^R^-GFP WT or mutant were measured in ciliated GFP-positive cells using Fiji version 1.54d (https://fiji.sc/). To calculate cilium length, the cilium was traced by a line, and the length of the line was measured in Fiji. The calculation of the *CPIR* was performed as described previously [27]. Briefly, first, a ∼1 μm thick line was drawn orthogonally across the cilium in Fiji. Next, the line intensity profile of the cilium was determined, and the maximum intensity value (*I*_max_) was obtained. Similarly, by drawing a circular ROI (region of interest) on the PM and the background region, the mean intensity of PM (*I*_PM_) and the background value (*I*_background_) were acquired, respectively. Finally, the CPIR of ARL13B^R^-GFP WT or mutant was calculated as (*I*_max_ − *I*_PM_)/(*I*_PM_ − *I*_background_).

Data analysis and graphs were plotted using OriginPro version 9.9 (OriginLab,https://www.originlab.com). The sample size n is indicated wherever applicable in figures or corresponding legends. Data are presented as the *mean* ± *standard* deviation (s.d.). The two-tailed unpaired *t*-test was conducted in Excel (Microsoft, https://www.microsoft.com/enus/microsoft-365/excel). A *p*-value ≤ 0.05 was considered statistically significant.

## Acknowledgments

We would like to thank the following Feng Zhang (Massachusetts Institute of Technology, USA) for sharing the DNA plasmid (pLenti-CRISPR v2) with us.

## Author contributions

L. L. and D. M. conceptualization; D. M., H. M. C., and L. L. methodology; D. M. and H. M. C. analysis, investigation, and validation; D. M. and L. L. writing; L. L. supervision, project administration, and funding acquisition.

## Data availability

The datasets generated during the current study are available in the GenBank repository [PV240348 to 359] and are included in Supplementary File 2. The datasets are available from the corresponding author on request.

## Supplementary Information

This article contains supporting information.

## Abbreviations

The abbreviations used are listed below

AA: amino acid
CC: coiled-coil region
*CPIR*: the cilium to plasma membrane intensity ratio
IDR: intrinsically disordered region
KO: knock out
NS: not significant
PAL: palmitoylation lipid anchorage site
PTCH1: patched1
PRR: proline-rich region
sgRNA: single guide RNA
SMO: smoothened
WT: wild type.

