## Supplemental Fig. 1 for "Knocking out *ARL13B* completely abolishes primary ciliogenesis in cell lines"

Supplementary Figure 1

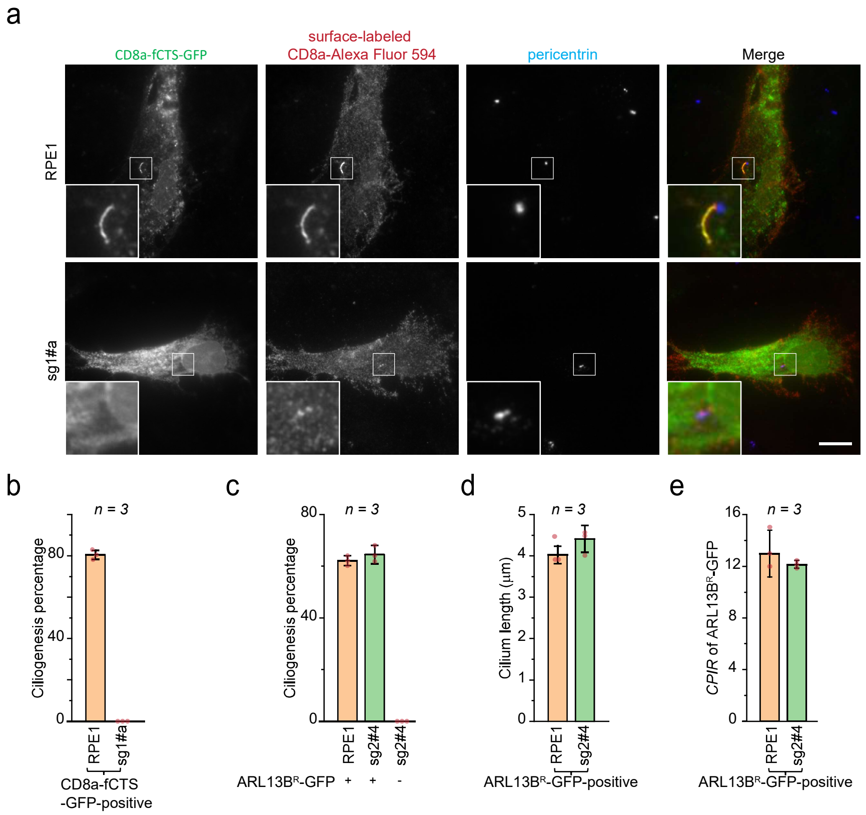


**Supplementary Figure 1**

*ARL13B*-KO RPE1 cell lines, sg1#a and sg2#4, have no cilia. (**a-b**) No cilia were observed in sg1#a cells using CD8a-fCTS-GFP as the cilium marker. Parental or sg1#a RPE1 cells expressing CD8a-fCTS-GFP were surfaced-labeled with anti-CD8a antibody before fixation and immunostaining for the CD8a antibody and the endogenous pericentrin (a). CD8a-fCTS-GFP positive cilia were only observed in parental but not sg1#a cells. Scale bar, 10 µm. The ciliogenesis percentage in CD8a-fCTS-GFP-positive cells was plotted in (b). (**c-e**) ARL13B$^{R}$-GFP expressing sg2#4 cells exhibit normal cilia. Images acquired as described in Figure 1g were analyzed. See the legend of Figure 1h-j.
