## Supplemental Fig. 2 for "Knocking out *ARL13B* completely abolishes primary ciliogenesis in cell lines"

Supplementary Figure 2

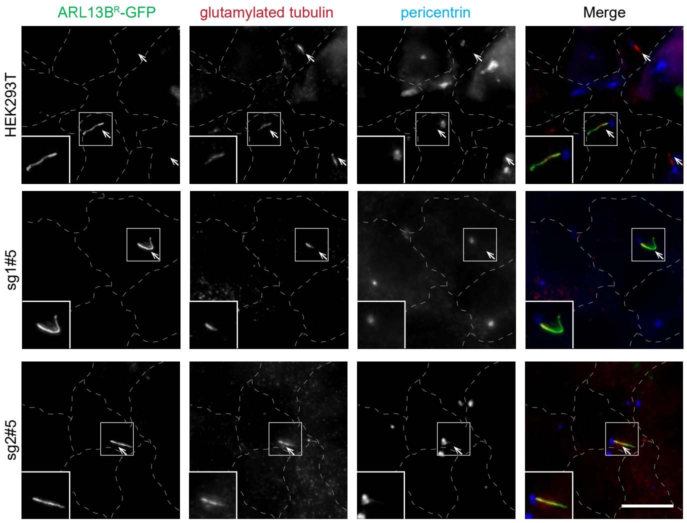


**Supplementary Figure 2**

ARL13B$^R$-GFP expressing HEK293T cell lines, sg1#5 and sg2#5, have normal cilia. (**a**) Indicated cells expressing ARL13B$^R$-GFP were fixed and processed for immunofluorescence labeling of endogenous glutamylated tubulin and pericentrin before being imaged by a wide-field microscope. Arrows indicate ciliated cells. Scale bar, 10 μm.
