## Supplemental Fig. 3 for "Knocking out *ARL13B* completely abolishes primary ciliogenesis in cell lines"

Supplementary Figure 3

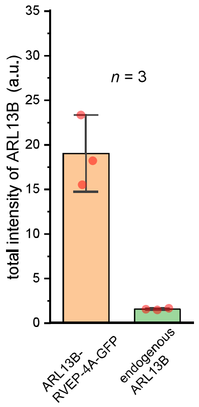


**Supplementary Figure 3**

Overexpression of transiently transfected ARL13B-RVEP-4A-GFP. Parental RPE1 cells, which transiently expressed ARL13B-RVEP-4A-GFP, were serum-starved to induce ciliogenesis and subsequently labeled with anti-ARL13B antibody. The total anti-ARL13B fluorescence was quantified and compared between GFP-positive (representing both ARL13B-RVEP-4A-GFP and endogenous ARL13B) and GFP-negative cells (representing only endogenous ARL13B). The data were obtained from $n = 3$ experiments, with more than 20 GFP-positive and GFP-negative cells counted in each experiment. The error bar represents the mean ± standard deviation.
