## Supplemental Fig. 4 for "Knocking out *ARL13B* completely abolishes primary ciliogenesis in cell lines"

Supplementary Figure 4

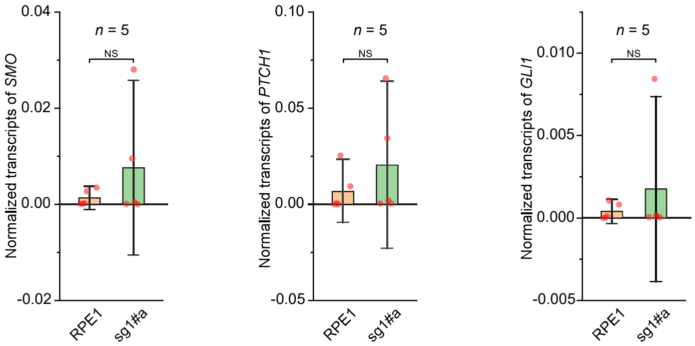


**Supplementary Figure 4**

Normalized transcription levels of *SMO*, *PTCH1*, and *GLI1* genes in parental or sg1#a RPE1 cells. The transcription was quantified by qPCR. *p*-values were from unpaired, two-tailed *t*-tests. NS, $p > 0.05$. $n = 5$ independent experiments.
