## Supplemental Fig. 5 for "Knocking out *ARL13B* completely abolishes primary ciliogenesis in cell lines"

Supplementary Figure 5

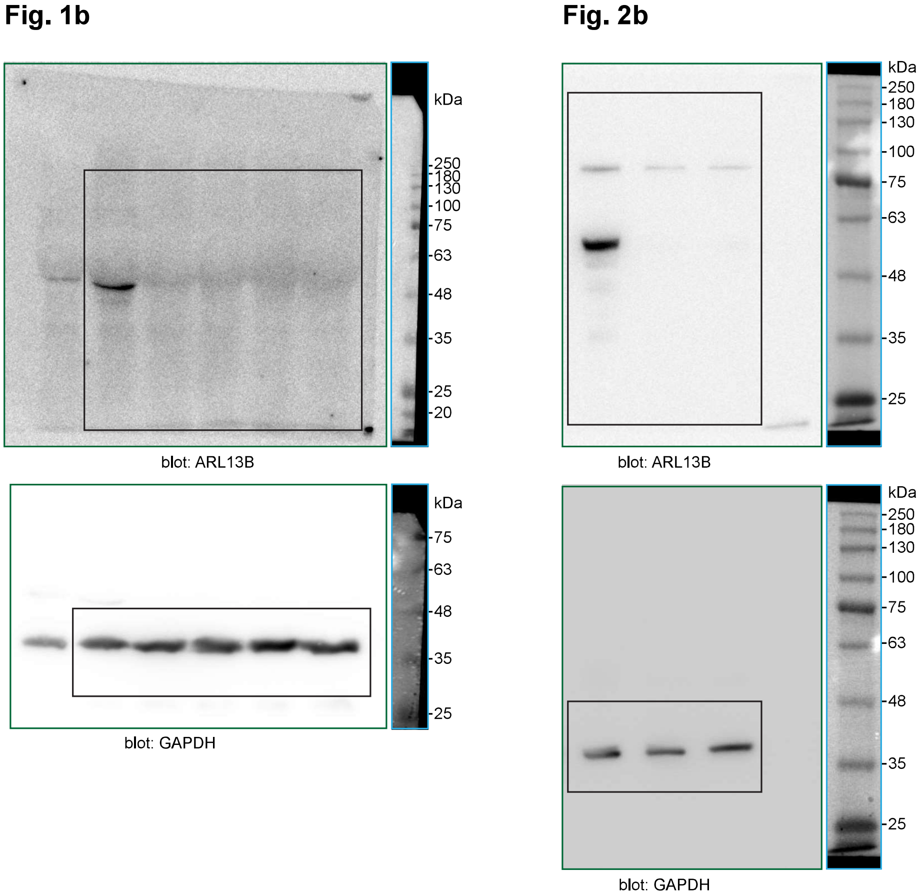


**Supplementary Figure 5**

Uncropped gel blot images used to prepare Fig. 1 and 2. Green box, chemiluminescence image; black box, cropped region that are shown in the corresponding figures; blue box, white light image of pre-stained molecular weight marker bands. Molecular weight (kDa) is labeled in all blots.
